# Nanoscale Structure, Interactions, and Dynamics of Centromere Nucleosomes

**DOI:** 10.1101/2024.02.12.579909

**Authors:** Shaun Filliaux, Zhiqiang Sun, Yuri L. Lyubchenko

## Abstract

Centromeres are specific segments of chromosomes responsible for the accurate chromosome segregation process. Centromeres are comprised of two types of nucleosomes: canonical nucleosomes containing an octamer of H2A, H2B, H3, and H4 histones, and CENP-A nucleosomes in which H3 is replaced with its analog CENP-A histone. This modification leads to the difference in the nuclear of DNA turns around the histone core, wrapping efficiency. This value is 121 bp of DNA, considerably less than 147 bp found in canonical nucleosomes. We used Atomic Force Microscopy (AFM) to characterize nanoscale features for both types of nucleosomes assembled on the same template, enabling us to evaluate the effect of internucleosomal interaction. We found that CENP-A mononucleosomes have a lower internucleosomal affinity than canonical H3 nucleosomes. We applied time-lapse, high-speed AFM (HS-AFM) to characterize the dynamics of nucleosomes. For both nucleosomes, spontaneous unwrapping of DNA was observed, and this process occurs via a transient state with ∼100 bp DNA wrapped around the core, followed by a rapid dissociation of DNA. The unwrapping process is asymmetric, so when the dissociation starts on one arm, it enlarges the size of the dissociated arm. Additionally, HS-AFM revealed higher stability of CENP-nucleosomes compared with H3 ones, in which dissociation of the histone core occurs prior to the nucleosome dissociation. The histone core of CENP-A nucleosomes remains intact even after the dissociation of DNA.

## Introduction

Nucleosomes are the fundamental nano assemblies in chromatin, the assembly of which is the first step for packing DNA in the nucleus.^1,2^ Interaction between nucleosomes is a fundamental property that defines the assembly and function of chromatin. Studies over the past two decades have revealed highly dynamic features of nucleosomes that can explain regulatory processes at the chromatin level (e.g., see recent reviews ^3–5^). However, structural details and the mechanism underlying the assembly of nucleosomes in higher-order structures of chromatin and their dynamics remain unexplained. Many cellular processes, such as transcription, require the dissociation of DNA from nucleosomes, which is achieved through nucleosome dynamics and remodeling machinery.^6,7^ Structural and single-molecule studies of these processes have been critical in developing current nucleosome models ^8^; however, the strong reliance on nucleosome positioning sequences for these techniques raises the question of how nucleosome structure and dynamics differ for those assembled on positioning vs. non-positioning DNA sequences.^5^ We used DNA templates with different sequences and AFM visualization to directly characterize the role of the DNA sequence on the positioning of nucleosomes and their interactions.^5,9–13^ In paper ^13^, we used DNA templates with different sequences and found that nucleosomes are capable of close positioning with no discernible space between them, even in the case of assembled dinucleosomes. This array morphology contrasts with that observed for arrays assembled with repeats of the nucleosome positioning motifs separated by uniform spacers.^14^ Simulated assembly of tetranucleosomes by random placement along the substrates revealed that the interaction of the nucleosomes promotes nucleosome array compaction.^13^ We developed in this paper a theoretical model capable of accounting for the role of DNA sequence and internucleosomal interactions in forming nucleosome structures. These findings suggest that, in the chromatin assembly, the affinity of the nucleosomes to the DNA sequence and the strengths of the internucleosomal interactions are the two major factors defining the compactness of the chromatin.

Two types of nucleosomes exist. Canonical nucleosomes (H3_nuc_) are found throughout the chromosome and consist of two of each histone (H2A, H2B, H3, and H4).^15–17^ The H2A and H2B form dimers and interact with the entry-exit site opposite the H3/H4 tetramer arranged at the dyad.^18^ The H3_nuc_ wrap ∼147 bp of DNA corresponding to ∼1.7 turns around the octameric histone core.^19–22^ A unique area of the chromosome is the centromere, which is responsible for holding together the sister chromatid and then must be pulled apart during replication.^23–25^ In the centromere nucleosomes, a variant of H3 histone is replaced with its variant CENP-A in the octameric core. CENP-A nucleosomes (CENP-A_nuc_) typically wrap 121 bp.^26,27^ Both types of nucleosomes are dynamic, and in our AFM experiments^9^, we found that CENP-A nucleosomes are capable of spontaneous unwrapping, which is the major dynamics pathway. ^9,28–30^ Unwrapped CENP-A nucleosomes can undergo long-range translocation by traveling over ∼200 bp; this process is also reversible.^28,29^ Additionally, CENP-A stabilizes nucleosome core particles against complete dissociation even when not fully wrapped with DNA.^28,29^

Here, we compared nanoscale features of both types of nucleosomes assembled on identical DNA templates. Using the DNA template with segments with different nucleosome affinity capable of forming two nucleosomes, we compared the interactions and dynamics properties of both types of nucleosomes assembled on the same DNA templates. Internucleosomal interaction was estimated by measuring the internucleosomal distance, revealing the elevated interactions between canonical nucleosomes compared with CENP-A ones. Time-lapse, high-speed AFM (HS-AFM) was applied to characterize the nucleosome’s unraveling dynamics, allowing us to reveal similarities and differences between canonical H3 and CENP-A nucleosomes.

## Experimental Methods

### Y-DNA construct

Our DNA preparation is done the same as our lab has done previously using PCR with a pUC57 plasmid vector from BioBasic (Markham, ON, CA).^10,31–33^ The 3WJ terminal DNA design was completed by introducing the SapI cut site in the primer (5’-TTAGCG**GAAGAGC**GCTTGTCTGTAAGCGGATGCCG-3’), with the cut site for SapI being bolded. The SapI cut site allows for a modular insertion of the restriction enzyme into any DNA construct. Once SapI cuts the DNA, construct a 3WJ ligated from three single-stranded DNAs, with the appropriate sticky end ligated to the original DNA. The full-length DNA containing the 3WJ end can be seen in Figure 1. There is a 601 motif that is 80 bp from the non-3WJ end. The rest of the DNA is made of a random sequence with no nucleosome binding specificity.

**Figure 1.**
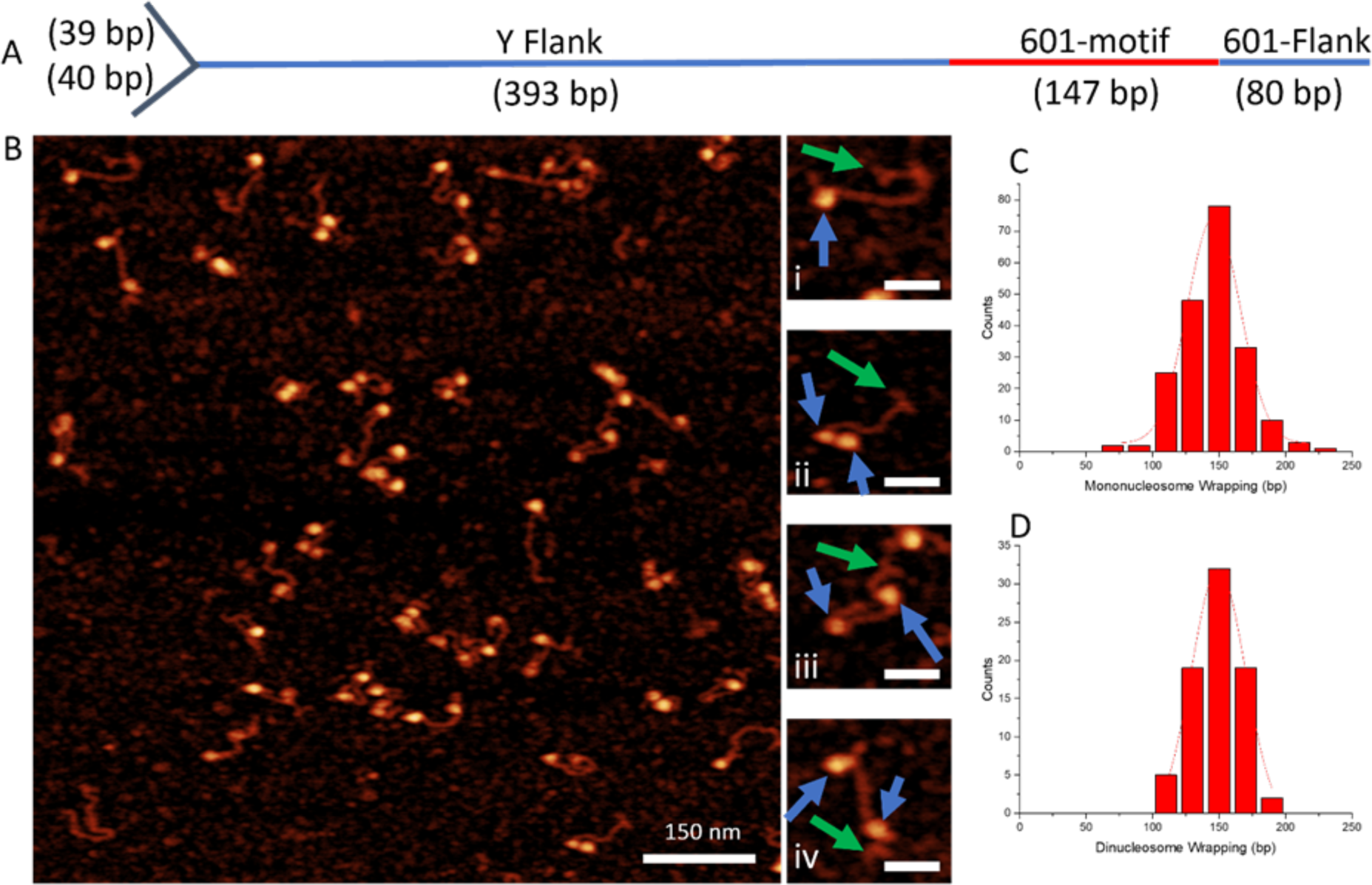
AFM imaging of H3 nucleosomes. – (A) DNA construct containing a three-way junction at one terminal end as a fiducial marker, and the 601 motif is shown in red. (B) AFM image 1000 nm x 1000 nm. Selected zoomed images of monoH3_nuc_(i) and diH3_nuc_(ii, iii, and iv) subsets are shown to the right of the main AFM image. Nucleosomes and 3WJ are indicated with blue and green arrows, respectively. (C, D) histograms for wrapping efficiencies of mononucleosomes (C) and the dinucleosomes (D).

### Nucleosome assembly H3 and CENPA

The nucleosome assemblies of both H3 and CENPA are completed through 24-hour dialysis, starting at 2 M NaCl and ending at 2.5 mM NaCl, the same as we have published previously.^28,30,31^ This process occurs at 4C and utilizes a peristaltic pump that continuously pumps the low salt buffer into the reaction beaker at the same rate that it pumps the high salt buffer out. The H3_nuc_ are purchased from The Histone Source (Fort Collins, CO) and are already in an octameric form. It requires mixing the nucleosomes and DNA and a predialysis to remove the glycerol from the stock. The CENP-A_nuc_ are purchased from EpiCypher (Durham, NC) and come in two stocks, one containing the H2A/H2B dimers and one containing the CENP-A/H4 tetramer. They require the extra step of mixing the two histone stocks in equimolar concentration to obtain an octameric core with two dimers and a single tetramer.

### Static AFM sample preparation and imaging

The static AFM samples are prepared after the assembly of the nucleosomes and are deposited on 1-(3-aminopropyl) silatrane (APS) mica, thereby functionalizing the surface of the mica. The stock solutions of nucleosomes (H3 and CENP-A) are stored at 300 nM at 4C. In preparation for the stock solution for imaging, a small aliquot is taken and diluted to 2 nM using imaging buffer (4 mM MgCl2 and 10 mM HEPES) and deposited on the APS mica. The mica containing the sample is incubated for 2 minutes, washed gently with DI water, and dried with a slow argon flow. The sample is placed in a vacuum and allowed to dry overnight under a vacuum. The dried samples are then imaged on a Multimode AFM/Nanoscope IIId utilizing TESPA probes (Bruker Nano Inc, Camarilla, Ca). The static images were captured at 3 x 3 um in size with 1536 pixels/line.

### High-Speed Atomic Force Microscopy Imaging in liquid

High-speed AFM imaging was performed as described in our previous literature.^28,30,34^ Briefly, a thin piece of mica was punched into 1.5 mm diameter circular pieces and then glued onto the sample stage of the HS-AFM (RIBM, Tsukuba, Japan). 2.5 μl of 500 μM APS solution was deposited onto the mica and incubated for 30 min in a wet chamber to functionalize the mica surface. The mica surface was then rinsed with 20 μl of deionized water. Then, 2.5 μl of the DNA or nucleosome sample was deposited onto the APS functionalized mica surface and incubated for 2 min. The sample was then rinsed with buffer and put into the fluid cell containing the imaging buffer described above. HS-AFM carried out imaging using electron beam deposition (EBD) tips. The typical scan size was 400 × 400 nm with an 800 ms/frame scan rate.

### Data Analysis

We utilize the same methods as previously published by our lab.^28,31^ The static images captured are analyzed using Femtoscan (Advance Technologies Center, Moscow, Russia), where we can measure the contour lengths of the DNA. The contour length measurements begin at the end of the 3WJ and are measured to the middle of the nucleosome. The second arm measurement starts at the center of the nucleosome and is measured to the 601 terminal end. 5 nm is subtracted from both arm lengths because of the contribution of the DNA to the wrapping around the nucleosome. A conversion factor is calculated from naked DNA to calculate the bp of the measurements. In the static images, measuring the contour lengths of all the free DNA and dividing by the known DNA length (659) will provide a number around 0.35, which we use to convert nm measurements to bp. The linker length between two nucleosomes is calculated by measuring the DNA length between the center of two nucleosomes and subtracting 10 nm of DNA to account for the contribution of both nucleosomes.

In HS-AFM, the contour length is calculated by measuring the DNA length after the nucleosome has evacuated the DNA, and the full-length DNA is accessible for measurement. The averages of the contour lengths are used to calculate the conversion factor. The histograms were created using Origin. Microsoft Excel was used for creating scatter plots and bar graphs.

## Results

### DNA substrate

We designed a DNA template capable of assembling two nucleosomes containing the nucleosome-specific 601 motif and non-nucleosome-specific random sequences to accomplish the goal. Schematically, the construct is shown in Figure 1A. At the end of the DNA, opposite to the location of the 601 motif, we placed a three-way junction DNA segment forming a Y-shape, which served as the marker for mapping the nucleosomes on the DNA.^12^ The nucleosomes were assembled as described in the methods section using a 2:1 molar ratio of the nucleosome core and DNA. The samples with CENP-A and canonical H3_nuc_ were assembled in parallel and prepared for AFM imaging as described earlier.^30,31^

### Positioning for canonical H3 nucleosomes

Figure 1B shows typical topographic AFM images for the array with canonical H3_nuc_ sample. Nucleosomes appear as bright globular features, and mononucleosome samples are seen along with dinucleosomes. Selected images for mono and dinucleosomes are shown to the right of the scan. The frame (i) shows a mononucleosome AFM image in which the nucleosome is indicated with a blue arrow, whereas the green arrow points to the Y-end of the DNA. Three other frames (ii) – (iv) illustrate dinucleosome samples with different distances between the nucleosomes indicated with blue arrows. The nucleosomes were found in close locations (frame (ii)) or far from each other, frames (iii) and (iv). The AFM images were analyzed to characterize the arrays.

First, the length of DNA wrapped around the nucleosome was measured to determine the length of DNA wrapped around the core, wrapping efficiency. It was done by subtracting the total contour lengths of DNA segments attached to the nucleosome core from the total length of the free DNA. The mono- and dinucleosome sample data are shown in Figure 1C and D, respectively. These data demonstrate that the monoH3_nuc_ wrap 145 ± 23 bp and the diH3_nuc_ wrap 149 ± 24 bp, which are in the expected 147 bp value range.^35,36^

Next, we mapped the position of nucleosomes for both types of samples. The data visualizing the positions of the center of the nucleosomes for the mononucleosome samples are shown in Figure 2A. The green highlighted area indicates the 601 location on the DNA construct. The zero position on the Y-axis corresponds to the DNA end opposite the 3WJ. The positions of the nucleosome center are marked with orange dots. The primary binding location of nucleosomes are to the 601 sequence (orange dots)—only three nucleosomes out of 202 (99%) bind to the locations outside the 601 region. The histogram of the nucleosome position in Figure 2B produces a narrow distribution.

**Figure 2.**
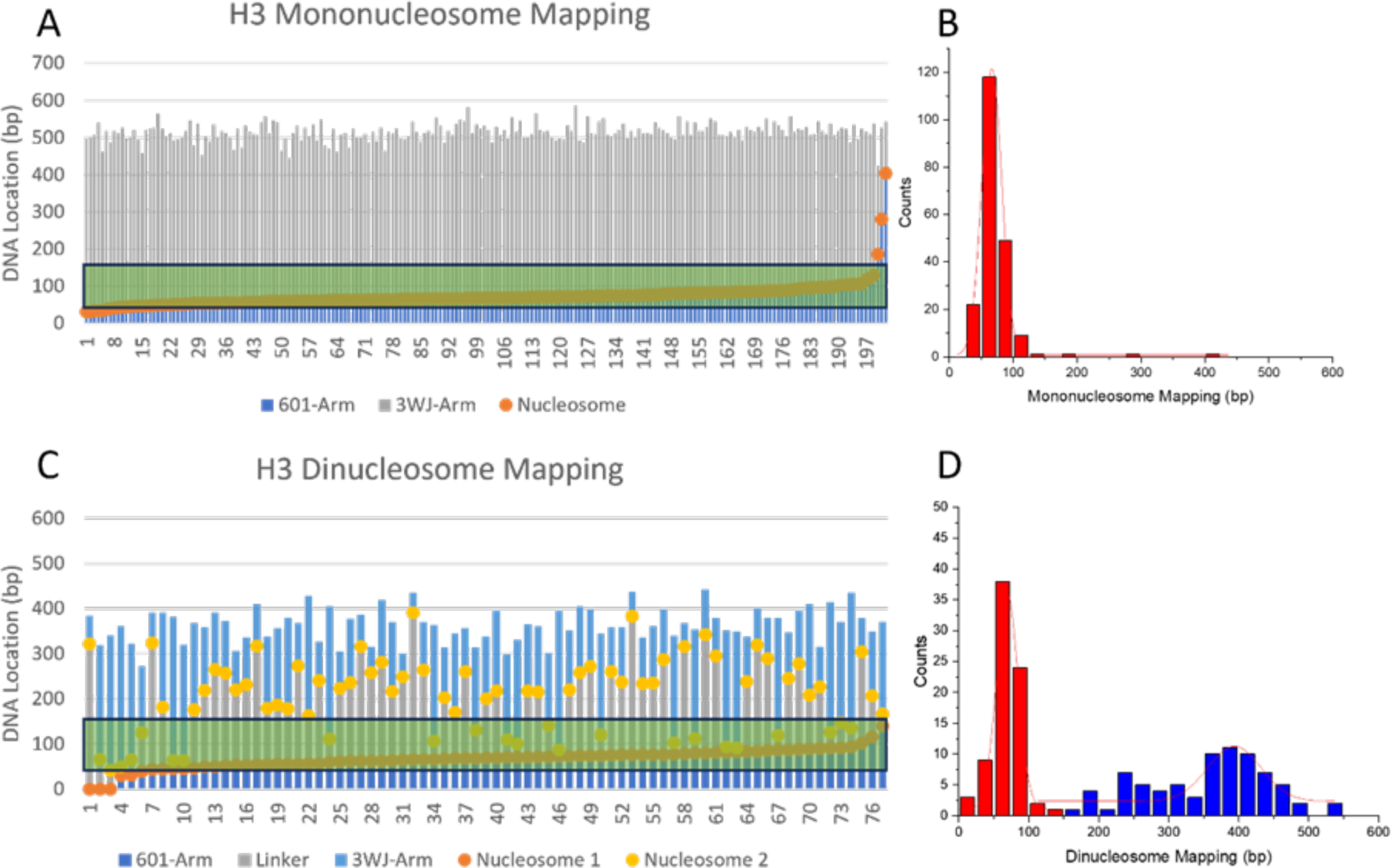
AFM mapping of H3_nuc_. The static AFM of monoH3_nuc_ (A) and diH3_nuc_ (B) mapping location binding on the DNA construct. The orange and yellow dots are nucleosome binding locations. The Y-axis is the DNA location, where 0 indicates the 601 terminal end and the 601 motif starts 80 bp from 0. The green area represents the 601 area on the DNA. (C, D) depict histograms for mapping data for mononucleosomes (C) and dinucleosomes (D). Different colors in C and D correspond to nucleosome position on 601 motif (red) and the rest of the DNA template (blue).

A similar mapping analysis was performed for the dinucleosome samples, and the results are assembled in Figure 2C. Nucleosomes bound to the 601 region are depicted in orange dots, and the position of the second nucleosome is shown as yellow dots. The positions of the yellow dots are not specific, so these are scattered over the rest of the DNA template. The histograms for the nucleosome positions assembled as histograms are shown in Figure 2D. A narrow peak (red) corresponds to the nucleosome assembled at the 601 sequence, and the positions of the second nucleosome shown in blue are not well defined, producing a broad peak.

### Positioning for CENP-A nucleosomes

Typical AFM images for the CENP-A_nuc_ samples are shown in Figure 3A with selected zoomed images of the subset’s mono- and dinucleosome species. In frame (i), a mononucleosome (blue arrow) can be seen bound to the 601 site, far from the 3WJ (green arrow) at the opposite end of the DNA. In frame (ii), two nucleosomes are relatively close to one another. In frame (iii), there is one stable nucleosome fully wrapped nucleosome near the 601 site, and near the 3WJ, there is an unwrapped nucleosome. In frame (iv), two partially unwrapped nucleosomes are bound to the DNA.

**Figure 3.**
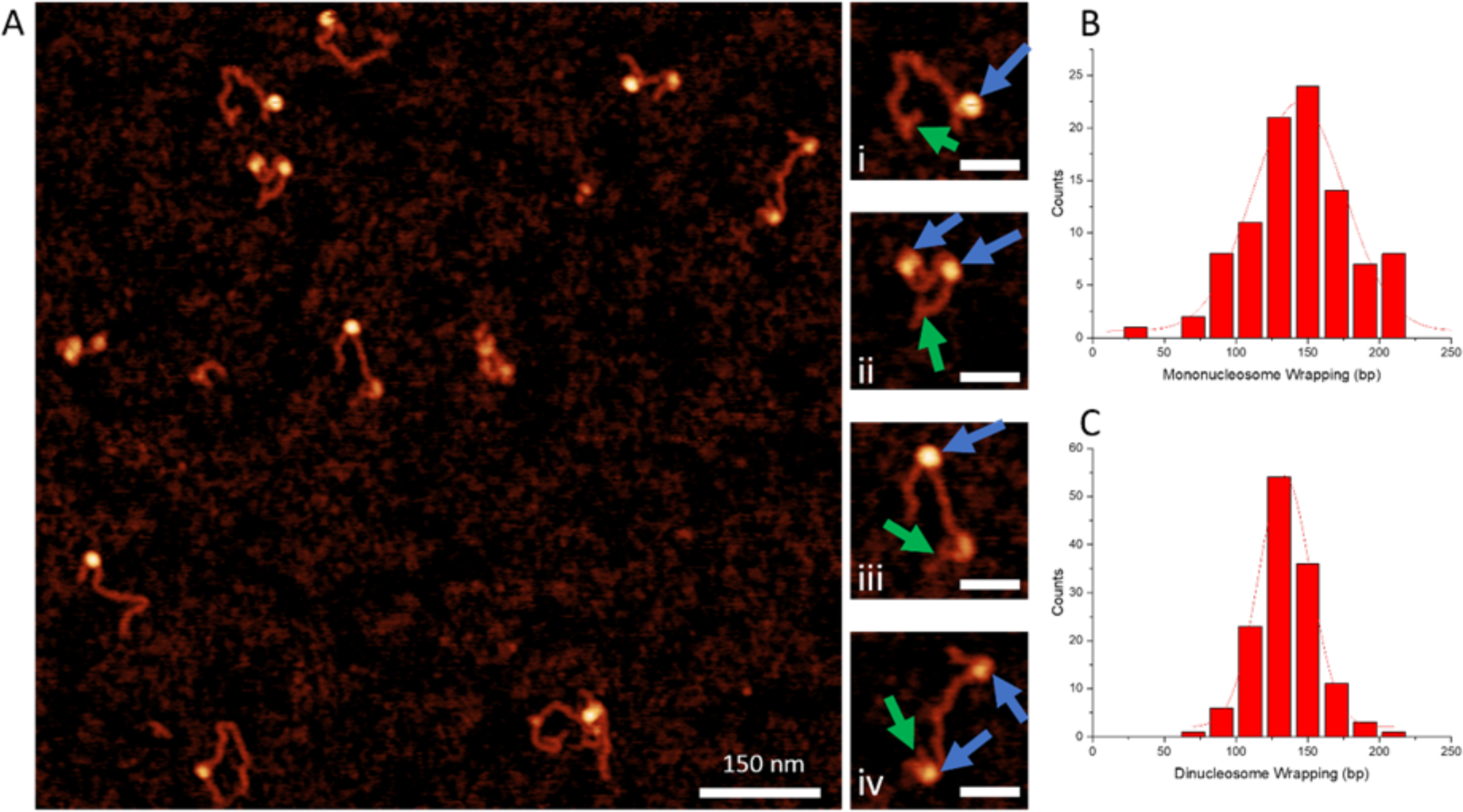
AFM imaging of CENP-A nucleosomes. (A) AFM image 1000 nm x 1000 nm. Selected zoomed images of monoCENP-A_nuc_ (i and iii) and diCENP-A_nuc_(ii and iv) subsets are shown to the right of the main AFM image. Nucleosomes and 3WJ are indicated with blue and green arrows, respectively. (B, C) histograms for wrapping efficiencies of mononucleosomes (B) and dinucleosomes (C).

The measurements of the DNA wrapping efficiency for CENP-A_nuc_ were done the same way as for the H3_nuc_ samples. The monoCENP-A_nuc_ samples had a DNA wrapping efficiency of 137 ± 43 bp; the standard deviation for the monoCENP-A_nuc_ wrapping is much larger than the H3_nuc_ counterpart. The dinucleosome assemblies of CENP-A_nuc_ DNA wrapping efficiency was 130 ± 20 bp. Histograms for the bp wrapping of mononucleosomes and dinucleosomes can be seen in Figures 3B and C.

The mapping of CENP-A_nuc_ was completed on the same DNA construct as the H3_nuc_. The monoCENP-A_nuc_ mapping results can be seen in Figure 4A.

**Figure 4.**
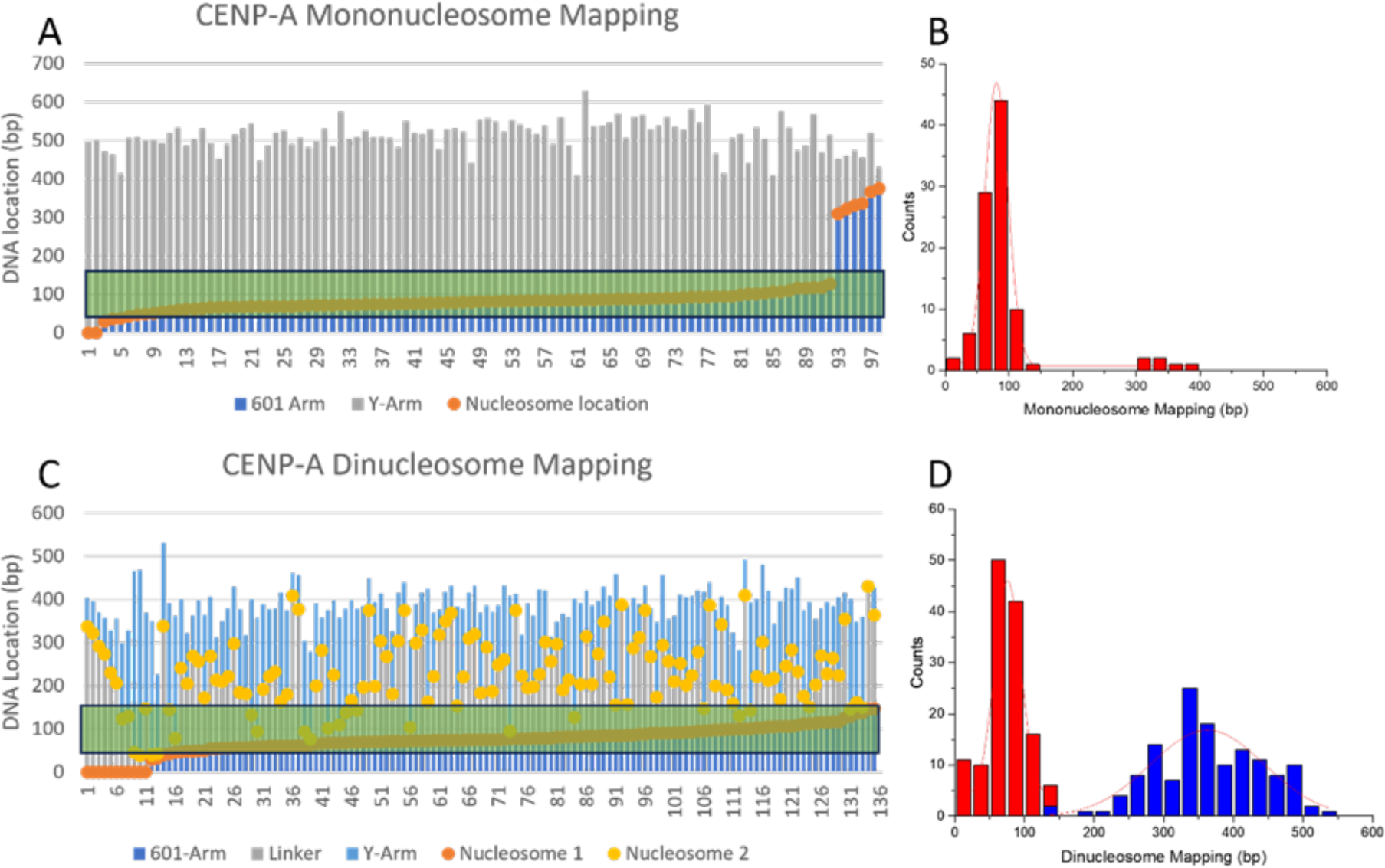
AFM mapping of CENP-A_nuc_. The static AFM of monoCENP-A_nuc_ (A) and diCENP-A_nuc_ (B) AFM mapping location binding on the DNA construct. The orange and yellow dots are nucleosome binding locations. The Y-axis is the DNA location, where 0 indicates the 601 terminal end and the 601 motif starts 80 bp from 0. The green area represents the 601 area on the DNA. (C, D) depict histograms for mapping data for mononucleosomes (C) and dinucleosomes (D). Different colors in C and D correspond to nucleosome position on 601 motif (red) and the rest of the DNA template (blue).

The mapping results for the dinucleosome CENP-A_nuc_ sample (diH3_nuc_) are shown in Figure 4C. The orange dots are the nucleosomes bound closer to the non-3WJ end, which results in 93% of the nucleosomes binding to the 601 sequence. The yellow dots represent the nucleosomes binding closer to the 3WJ end, which is comprised of non-specific DNA; therefore, the nucleosomes have random binding locations. Interestingly, there is an elevated affinity of the CENP-A_nuc_ assembly at the end of the DNA template. The green highlighted area shows the location of the 601 sequence on the DNA construct. The orange dots represent the center of the nucleosome binding location. The blue arm represents the DNA from the center of the nucleosome to the non-3WJ terminal end, and the grey bars represent the DNA from the center of the nucleosome to the 3WJ terminal end. The high affinity of the nucleosomes to the 601 motif is seen in these nucleosomes, although there is a decrease to 92% binding to the specific sequence as compared to 99% for H3_nuc_. A histogram representation of the CENP-A_nuc_ binding location can be seen in Figure 4B, showing the specific binding to the 601 region.

### Internucleosomal interactions for H3 and CENP-A nucleosomes

Nucleosomes in the AFM images (e.g., plate (ii) in Figure 1B) are located close to each other, pointing to the interaction between the nucleosomes. Such events can be identified in dinucleosome maps (Figure 5) by the co-localization of two nucleosomes. We used the values for the nucleosome locations’ centers to measure the linker length’s internucleosomal distance to characterize the ratio of such close contacts.^10^ The results of such measurements for canonical H3 dinucleosomes are shown as histograms in Figure 5A. The first bar in the histogram corresponds to the close location of the nucleosome, which, according to our publication ^10^, points to the formation of close internucleosomal contacts. The yield of nucleosomes with a linker length of less than 50 bp was 23%. A similar analysis was done for the CENP-A dinucleosomes, and the histogram is shown in Figure 5B. Although the bar corresponding to the distance below 50 nm is detectable, its height is twice lower than that for H3_nuc_ (Figure 5A), pointing to the weaker internucleosomal interactions for CENP-A_nuc_ compared with H3_nuc_ ones.

**Figure 5.**
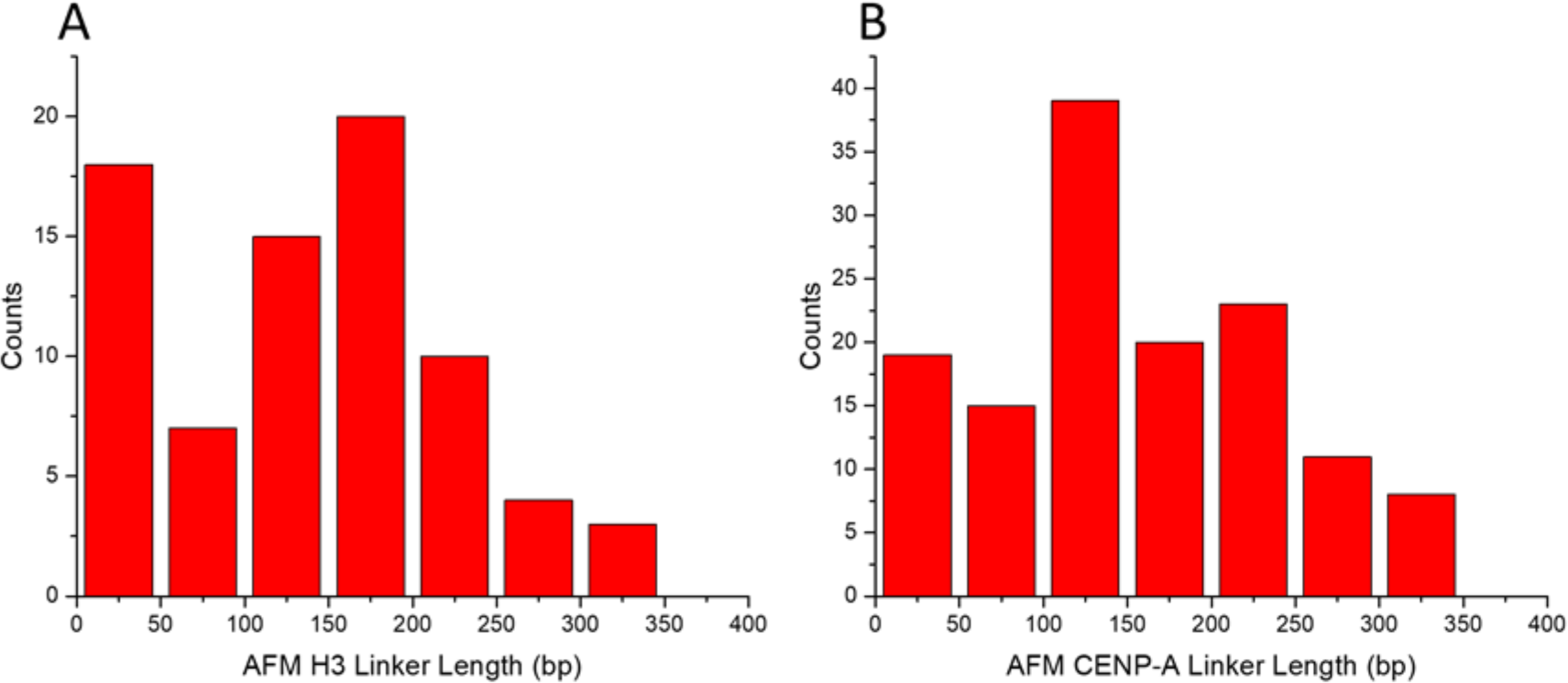
Dinucleosome linker length results. The H3_nuc_ linker length (A) shows a preferential linker length of less than 50 bp and a gaussian distribution of around 170 bp. The CENP-A_nuc_ linker length (B) demonstrated a lower yield of linker lengths less than 50 bp and the largest population of around 125 bp.

### Time-lapse AFM to Probe Nucleosome Dynamics

Nucleosomes are dynamic complexes capable of spontaneous dissociation, which were directly characterized by single-molecule approaches.^37,38^ AFM is attractive among these methods because it can directly visualize the spontaneous unraveling of nucleosomes using the time-lapse AFM approach.^38–40^ In this approach, the sample is placed on the substrate, and continuous scanning over the same area allows one to observe the dynamics of various systems, including nucleosomes.^40^ We applied high-speed AFM capable of data acquisition in the millisecond time scale ^34,41^ to compare the dynamics of two types of nucleosomes characterized above.

Multiple frames are acquired, and a set of time-lapse images over the same area is shown in Figure 6 to illustrate a spontaneous unraveling of the nucleosome—the complete set of datasets with 211 frames assembled as Movie S1 is shown in the supplement. There were approximately 100 videos analyzed in total for H3_nuc_ and CENP-A_nuc_, which resulted in 4113 total frames analyzed.

**Figure 6.**
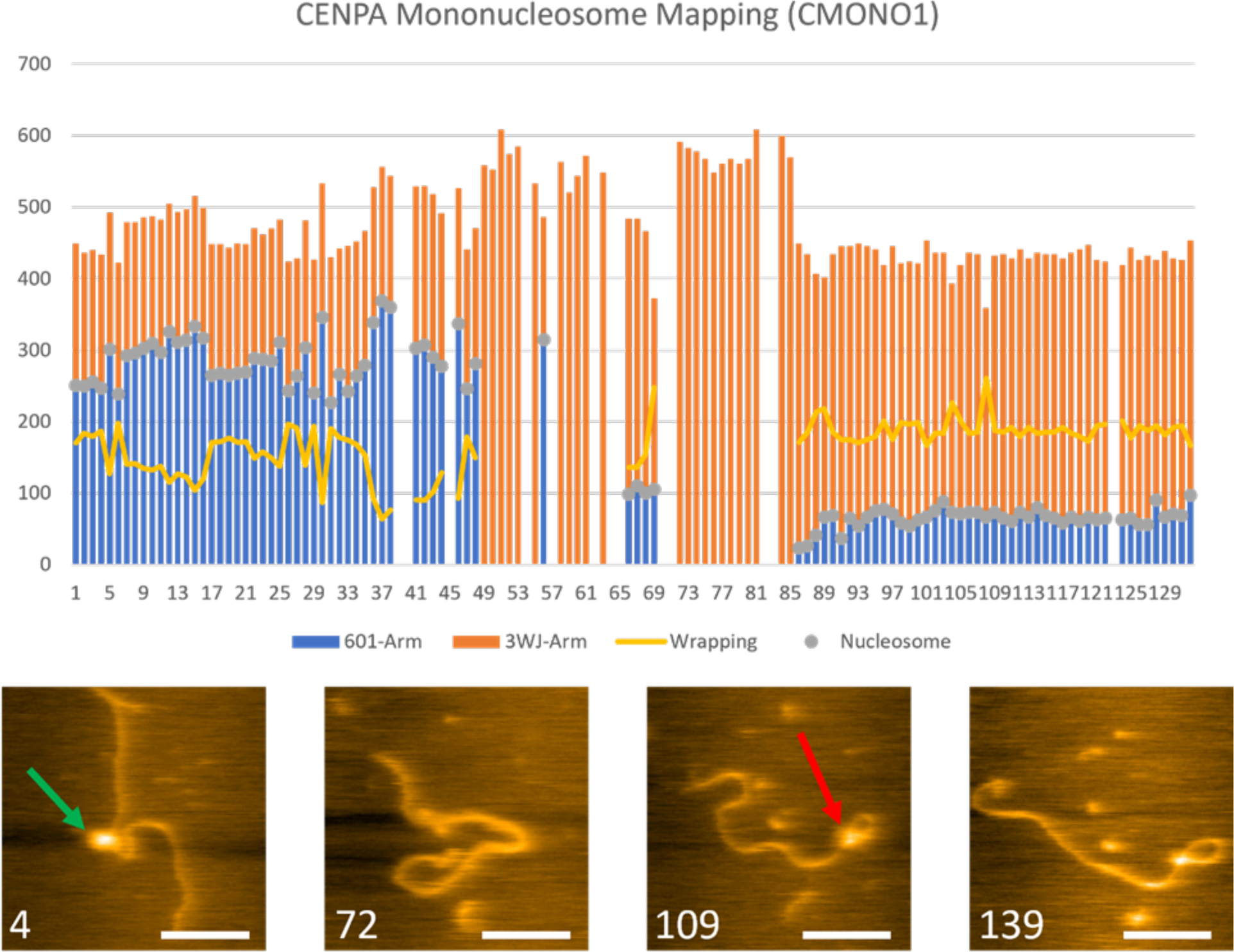
High-speed AFM video analysis. The Movie S1 analyzed, showing the mapping (bar graph) and snap shots of particular frames. The grey dots in the bar graph indicate the nucleosome bound to the DNA. The yellow line shows the wrapping of the CENP-A nucleosome. The number in white in the bottom left of the snap shots indicates the frame the image is from. In frame 4, the nucleosome is fully wrapped in a random location (green arrow) in the middle of the DNA. In frame 72, the nucleosome has disassociated completely from the DNA. In frame 109, a nucleosome rewraps DNA, spontaneous assembling a nucleosome. This assembly remains stably wrapped up to frame 139. Scale bar is 50 nm.

### Dwell time for nucleosomes unraveling

One of the parameters we analyzed was the dwell time of the nucleosome, defined by the time required for the complete unraveling of the nucleosome. First, we compared the dwell times of mono and dinucleosomes, and the results for H3_nuc_ can be seen in Figures 7A and B, respectively. For monoH3_nuc_, we found that 38% of the videos analyzed lasted 20 or more frames. This dwell time was increased by 48% for the diH3_nuc_, suggesting the nucleosome stability increased by the internucleosome interactions.

**Figure 7.**
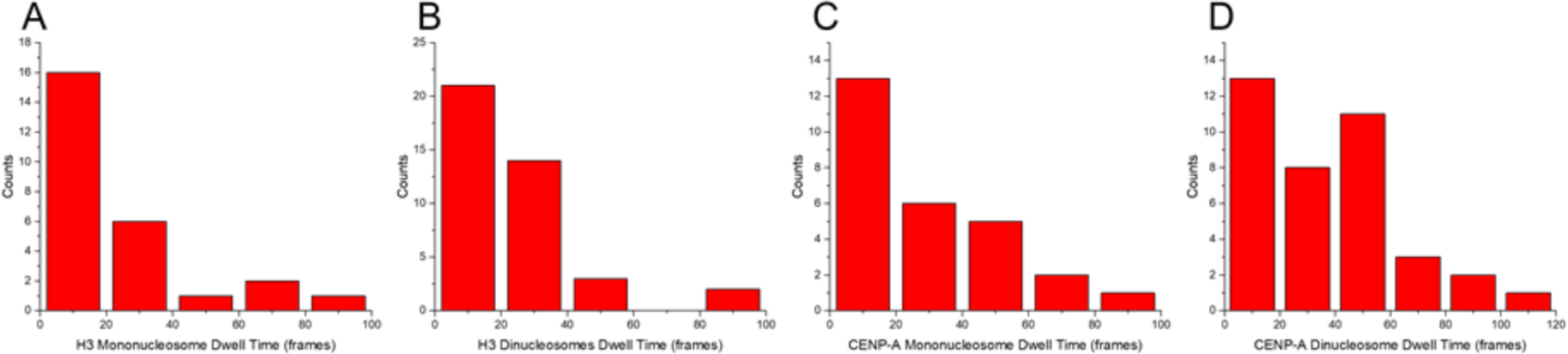
Nucleosome dwell times on HS-AFM. The monoH3_nuc_ results (A) showed most nucleosomes lasting 20 frames or less. The diH3_nuc_ results (B) showed a shift to the right, indicating that the dinucleosomes have a longer dwell time than the mononucleosomes. The monoCENP-A_nuc_ results (C) showed that most nucleosomes lasted less than 20 frames. The diCENP-A_nuc_ results (D) showed a shift to the right, indicating that the dinucleosomes have a longer dwell time than the mononucleosomes.

A similar analysis was done for CENP-A_nuc_ samples, and the data are summarized as histograms in Figure 7C and D. For the monoCENP-A_nuc_, 52% of the videos analyzed lasted longer than 20 frames, and 66% for the dinucleosomes. CENP-A_nuc_ followed a similar trend as with H3_nuc_, but this time, an increase of 14% from mononucleosomes to dinucleosomes. Also of interest is that the CENP-A_nuc_ averaged longer dwell times than the H3_nuc_, for monoH3_nuc_ with 38% and CENP-A_nuc_ with 52% of videos with longer than 20 frames in a video. The difference between H3_nuc_ and CENP-A_nuc_ is 14%. The same trend is visible for the dinucleosome results, with a difference of 18% more diCENP-A_nuc_ lasting longer than 20 frames than H3_nuc_. According to these studies, CENP-A_nuc_ are more stable than H3_nuc_ nucleosomes.

### Dynamics of nucleosomes core during unraveling

This large set of data revealed that unraveling is a non-gradual process. It is illustrated in Figure 8. One set of images is shown in Figure 8A (Movie S2). It can be seen that there is an H3_nuc_ (green arrow) that is wrapped at the 601 location (frame A1). In the following frames (A2 and A20), the histone (blue arrow) has vacated the octameric core, but the partial core remains to wrap the DNA. In frame A36, the DNA unwraps the partial core, resulting in a wrapping of ∼100 bp, and the free histone remains near the DNA filament. In the last frame, A66, the DNA loosens even more, effectively no longer tightly wrapping DNA around the core. Another example of a histone exiting the octameric core can be seen in Figure 8B (Movie S3).

**Figure 8.**
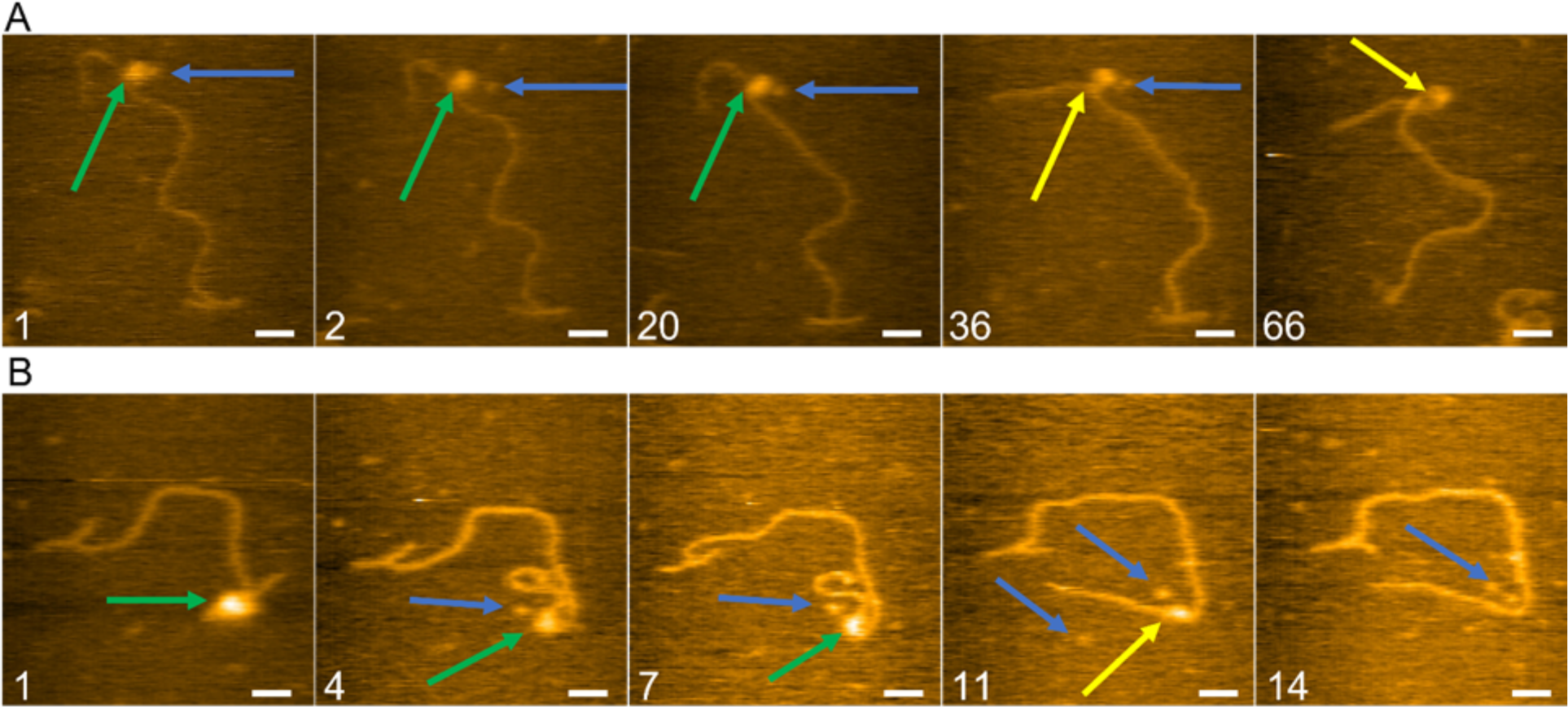
HS-AFM analysis of H3_nuc_ unwrapping. HS-AFM videos analyzed, showing the protein unwrapping pathway of H3_nuc_(green arrows) with histones (blue arrows) leaving the nucleosome core particle. Once the DNA unwraps the non-octameric core, they are indicated with yellow arrows—the frames in A come from Movie S2. In frame, A1, the nucleosome is bound to the 601 location, and the histone can be seen bulging out, indicated with the blue arrows. This histone moves in frames A2 and A20 while the DNA and nucleosomes remain in the same location. In frame A36, the DNA unravels the nucleosome; by frame, A66, only the tetramer can be seen still bound to the DNA. The frames in B come from the video of Movie S3. In frame, B1, the green arrow indicates a fully wrapped nucleosome bound to the 601 location. In frames B4 and B7, the histone (green arrow) can be seen to have left the nucleosome core particle, leaving a partial core. By frame B11, the DNA has unwrapped the partial core, and the tetramer remains bound. By frame B14, all histones have evacuated the DNA. The scale bar represents 25 nm.

Next, we looked at the unwrapping pathways of CENP-A_nuc_ on the same DNA construct. Figure 9A (complete set of frames in Movie S6), frame A1 shows a fully wrapped CENP-A_nuc_ (green arrow) bound to the 601 site. The nucleosome remains stably wrapped in frames A4 to A18. In frame A20, the nucleosome (yellow arrow) spontaneously unwraps, resulting in a decreased wrapping of ∼110 bp. The CENP-A_nuc_ remains wrapped ∼110 bp through the next frame shown, A37. The example in Figure 9B (Movie S7) shows a CENP-A_nuc_ bound near the 3WJ (frame B3). The CENP-A_nuc_ (green arrow) remains stably wrapped at this location through frames B8 to B11. In frame B13, the nucleosome (yellow arrow) undergoes unwrapping, resulting in a wrapping of ∼100 bp. By frame B18, the nucleosome no longer wraps any DNA. Notably, no octamer dissociation is observed during the entire unraveling process.

**Figure 9.**
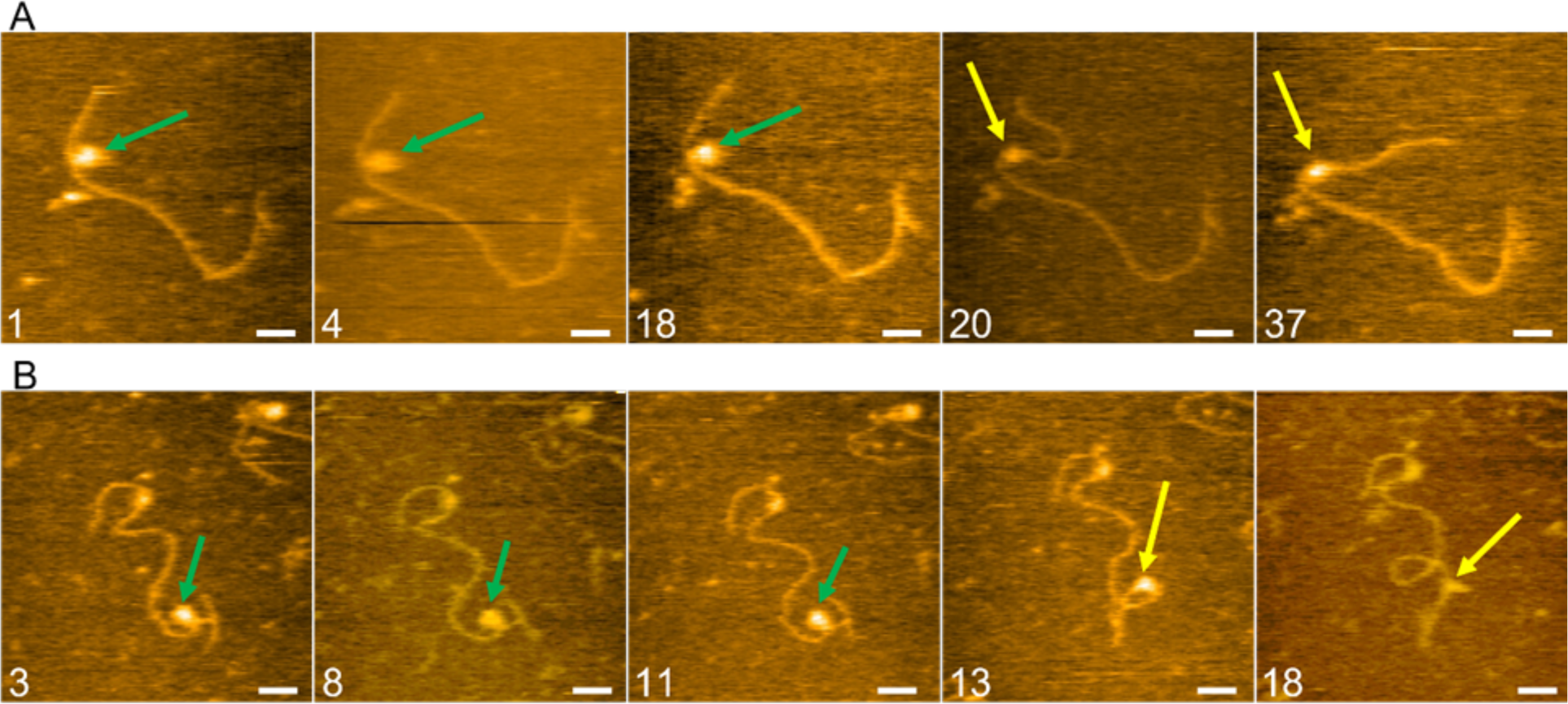
HS-AFM analysis of CENP-A_nuc_ unwrapping. AFM videos were analyzed, showing the DNA unwrapping pathway of CENP-A_nuc_ (green arrows). Once the DNA unwraps the nucleosome core particle, they are indicated with yellow arrows—the frames in A come from Movie S6. In frame A1, the nucleosome can be wrapped fully at the 601 location. The nucleosome remains fully wrapped in frames A4 and A18. Suddenly in frame A20, the DNA unwraps, leaving the nucleosome core partially wrapped (∼110 bp), indicated by the yellow arrow. The nucleosome particle remains partially even into frame A37 (∼100 bp). The frames in B come from the video of Movie S7. In frame, B3, the green arrow indicates a fully wrapped nucleosome bound near the 3WJ. In frames, B8 and B11, the nucleosome (green arrow) is still fully wrapped. By frame B13, the DNA has unwrapped the nucleosome (∼100 bp). By frame B18, the nucleosome no longer wraps any DNA. The scale bar represents 25 nm.

The nucleosome core stability of H3_nuc_ was found to be weaker through our analysis of the HS-AFM videos. We found that out of 69 total movies of H3_nuc_ analyzed, 53 (77%) of them resulted in the nucleosome core losing histones during the unwrapping, as indicated in Figure 5A and B, the blue arrows are pointing to the histones leaving the nucleosome core. In contrast, CENP-A_nuc_ had 63 nucleosomes analyzed, and only 6 (10%) had histones leave the core during the unwrapping event, indicating that the DNA was able to unwrap the nucleosome core without the octameric core falling apart. These results lead us to believe that the CENP-A_nuc_ core particle is significantly more stable than the H3_nuc_, which likely plays a role in its necessity of having a strong interaction to withstand the kinetochore pulling the sister chromatid apart. This overall higher stability of CENP-A_nuc_ is also supported by the longer dwell times found with CENP-A_nuc_ compared to the H3_nuc_, as seen in Figure 7.

### A step-wise nucleosomes unraveling

AFM images shown above point to the step-wise process of the nucleosome unraveling. To characterize this phenomenon, we measured the lengths of the DNA arms for each frame and plotted these measurements as a set of bars with different colors. These data are shown in Figure 10A, and a few snapshots are displayed in Figure 10B.

**Figure 10.**
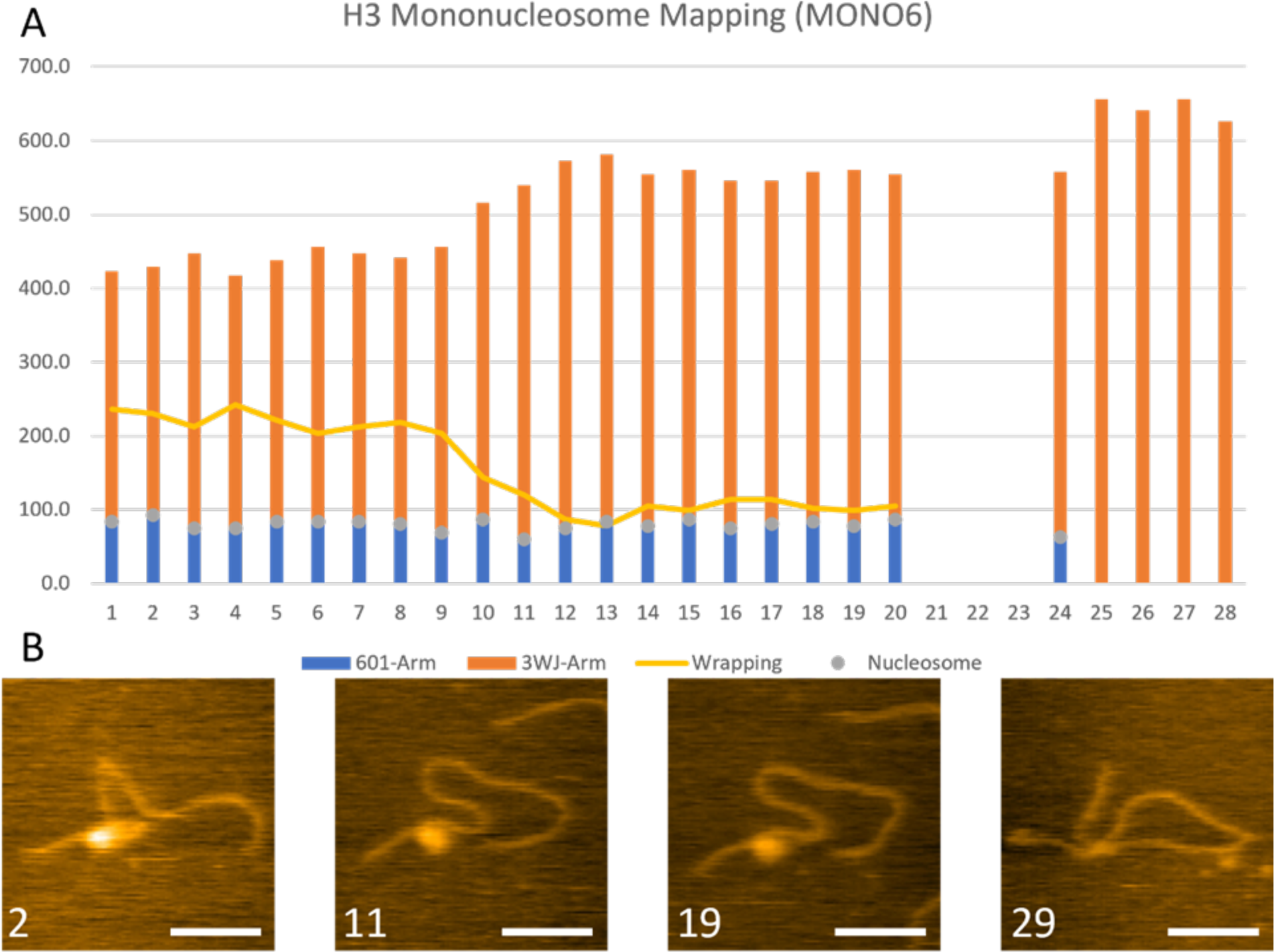
HS-AFM video of H3_nuc_ analyzed with snapshots. - The Movie S4 was analyzed, showing the mapping (bar graph) and snapshots of particular frames. The orange bars represent the DNA from the 3WJ end to the center of the nucleosome (grey dot). The blue bar represents the 601 DNA, from the 601 end to the center of the nucleosome. The yellow line shows the wrapping of the H3_nuc_. The number in white in the bottom left of the snapshots indicates the frame of the image.

The complete set of frames are assembled into Movie S4. The blue bars represent the distance from the 601 end of the DNA to the center of the nucleosome (grey dot). The orange bar represents the distance from the center of the nucleosome to the end of the 3WJ. The yellow line shows the DNA wrapping values calculated from the length measurements of the arms. In this set of images in the frame (2), the nucleosome can be seen to be overwrapped. These data show that there was a partial unwrapping event that started in frame (11) and proceeded to frame (13), where the nucleosome went from a state of overwrapped (∼210 bp) and, throughout three frames, it decreased to an under wrapped state (∼100 bp), where it remained bound for another 7 frames. The last frame 29 shows the nucleosome wholly disassociated from the DNA.

A similar event is shown in the supplementary Figure S1 and in the Movie S5. It was demonstrated that the initial overwrapped state (∼200 bp) in frame 10 dropped quickly to an under-wrapped state (∼100 bp), where it remained for another 6 frames.

Both videos demonstrated an intermediate step in H3_nuc_ disassembly, a state in which the nucleosome is considerably stable. There was a variation in the time that these unwrapping events took place. In Movie S4, the unwrapping took place over 3 frames (2.4 s), whereas in Movie S5, the unwrapping took place over 1 frame (0.8 s). The difference in time to unwrap indicates there is still something not completely understood in this process.

A similar analysis was done on CENP-A nucleosomes. The data are shown in Figure 11, where the results of measurements are shown (Figure 11A), and snapshots are displayed below (Figure 11B). The complete set of frames are assembled into Movie S8. In the snapshots shown, frames (4) to (36) show a fully wrapped nucleosome, and frame (43) shows an increase in DNA length and a decrease in bp wrapping around the nucleosome. Finally, frame (79) shows histones bound to the DNA without wrapping. According to Figure 11A, the CENP-A_nuc_ was stability-wrapped (∼130 bp) for 41 frames, at which point it went to an unwrapped state (∼90 bp). The unwrapping process took only a single frame (0.8 s).

**Figure 11.**
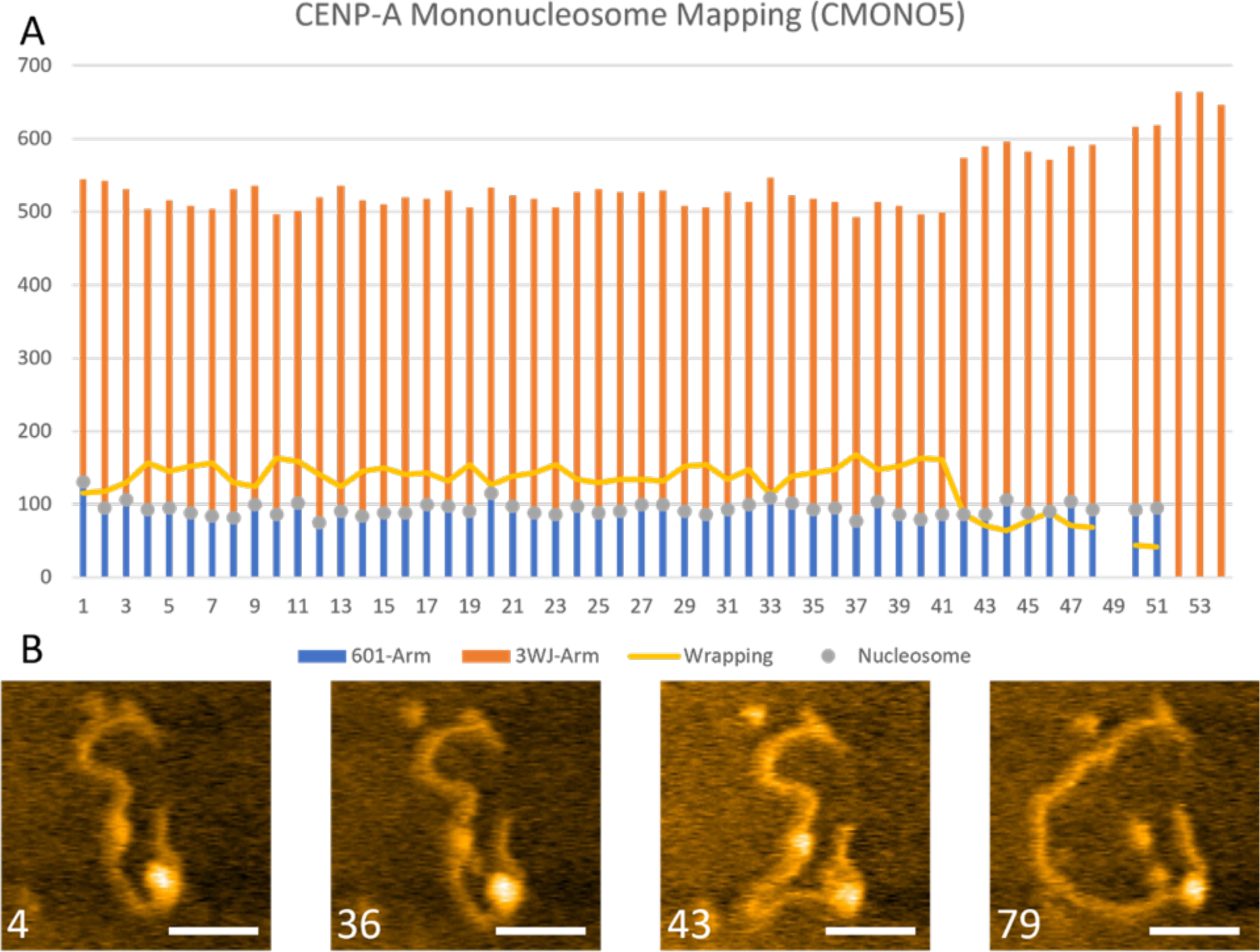
HS-AFM video of CENP-A_nuc_ analyzed with snapshots. The Movie S8 was analyzed, showing the mapping (bar graph) and snapshots of particular frames. The grey dots in the bar graph indicate the nucleosome bound to the DNA. The orange bars represent the DNA from the 3WJ end to the center of the nucleosome (grey dot). The blue bar represents the 601 DNA, from the 601 end to the center of the nucleosome. The yellow line shows the wrapping of the CENP-A_nuc_. The number in white in the bottom left of the snapshots indicates the frame of the image.

Another video with the same analysis was done to confirm the step-wise disassembly process of the previous video, seen in supplementary Figure S2. This video demonstrates a similar event as Figure 10. The nucleosome began in an over-wrapped state (∼200 bp), remaining for 33 frames. The nucleosome was unwrapped from frame 33 to 35 to ∼100 bp, staying for another 12 frames before completely disassociating.

### Asymmetric unraveling of nucleosomes

A closer analysis of the H3_nuc_ (Figs. 10 and S1) and CENP-A_nuc_ (Figs. 11 and S2) movies revealed that the unwrapping process is asymmetric, so one arm increases the size without changing another arm in length. In Figure 10A, the H3_nuc_ short 601 arm (blue) remains constant throughout the video, but the 3WJ arm (orange) increases in length during the unwrapping. Similar asymmetry was observed for the nucleosome initially assembled away from the 601 motif (Figure S1). The blue arm fluctuated, but the orange bar grew as unwrapping occurred (yellow line in Figure S1A).

The asymmetry in the unwrapping was observed for CENP-A nucleosomes, which is illustrated in Figure 11. The nucleosome is bound to the 601 site, and the short arm (blue) remained constant throughout the video, but the 3WJ arm (orange) grew in length during the unwrapping. In supplementary Figure S2, the CENP-A_nuc_ was assembled near the middle of the DNA template, away from the 601 motif. The nucleosome length of both arms remained relatively consistent until the unwrapping event, which caused the blue arm to grow in length with the orange bar remaining constant. Thus, the asymmetric unwrapping events do not have a preference for the DNA sequence.

In the analysis of 98 total unwrapping events, asymmetric unwrapping was found in 87% of cases for H3_nuc_ and 88% of the CENP-A_nuc_. The symmetric unwrapping was observed in 13% and 12% for H3_nuc_ and CENP-A_nuc_, respectively.

## Discussion

The dinucleosome approach using end-labeled DNA revealed several different features of CENP-A_nuc_ and bulk H3_nuc_. Although both nucleosome types can form tight contact with no visible space between nucleosomes (Figure 1B, frame (ii)), CENP-A_nuc_ indicated a lower effect than H3_nuc_ ones (Figure 5). We have demonstrated that the balance between energies of internucleosomal interaction and the affinity of the nucleosome core to the DNA sequence previously defines the formation of tight contacts between the nucleosomes.^10,13^ The balance favors the dinucleosome assembly for non-specific DNA sequences. We have shown that the number of such contacts increases in nucleosomes assembled with truncated H4 histone, suggesting that histone tails contribute to the tightening of dinucleosomes.^10^ Therefore, we hypothesize that the lower yield of dinucleosome complexes for CENP-A_nuc_ can be due to the repulsion generated by the CENP-A tail, and we plan experiments to test this hypothesis.

The DNA sequence is the primary factor defining the nucleosome positioning in the chromatin, and the 601 motif is the strongest nucleosome positioning sequence—our AFM data in Figures 1B and 3A visually support it, illustrating the almost exclusive formation of mononucleosomes on the position of the 601 motif. At the same time, there is a difference between both types of nucleosomes. The yield of H3_nuc_ bound to the 601 site in the mononucleosome and dinucleosome samples was 99% (Figs. 2B and D). The CENP-A_nuc_ were mapped similarly to H3_nuc_ and found bound to 601 site at 92% for mononucleosomes and 93% for the dinucleosomes (Figs. 4B and D). These differences in the binding affinity to 601, although marginal ∼6%, were consistent in both the mono and dinucleosomes results. Although the DNA sequence and specifically the TA dinucleotides provide such a high affinity of 601 to the formation of nucleosomes, it was shown in^42^ that interaction with histones contributes to the nucleosome positioning and H3–H4 tetramer dominates in the DNA sequence dependency effect. Replacement of H3 histone with CENP-A histone can decrease this DNA affinity effect.

The elevated affinity of CENP-A_nuc_ to the DNA ends with another property of these nucleosomes, illustrated in Figure 3. DNA wrapping around CENP-A_nuc_ is less than for H3_nuc_, 137 bp +/-20 bp vs. 145 +/- 23 bp, respectively, which is consistent with previous publications.^11^

Time-lapse HS-AFM studies revealed different stabilities of H3 and CENP-A nucleosomes. The data in Figure 7 demonstrate that CENP-A_nuc_ appear more stable under the scanning conditions than H3_nuc_. In the dwell time analysis on the HS-AFM, we found that 38% of monoH3_nuc_ lasted longer than 20 frames, whereas 52% of monoCENP-A_nuc_ lasted longer than 20 frames, a substantial increase of 14%. The dinucleosomes dwell comparison also showed that H3_nuc_ had a dwell time of more than 20 frames only 48% of the time compared to the CENP-A_nuc_ at 66%, a dramatic increase of 18%. This finding seems counterintuitive as the wrapping efficiency of CENP-A_nuc_ is less than H3_nuc_. However, CENP-A nucleosomes are not simply partially unwrapped H3 nucleosomes. Structures of CENP-A and H3 nucleosomes are different, pointing to different contacts between the DNA and histones, so this structural property of these two types of nucleosomes explains their different stabilities. For example, the CENP-A nucleosomes have increased flexibility of the DNA ends, the octameric core is more rigid, and has a different surface charge (positive) at the interface of the L1 of CENP-A and L2 of H4; in contrast, this surface is negatively charged in H3_nuc_.^27,43^

Most commonly, the H3_nuc_ found in the unwrapped state also appeared to have lost some of the histones from the nucleosome core, an H2A/H2B dimer. The loss of the dimeric histone caused the nucleosome core to become unable to maintain the fully wrapped ∼147 bp and loosen to a state of ∼100 bp wrapping. The remaining dimer sits at the entry-exit site opposite the tetrameric H3/H4 dimer at the nucleosome’s dyad. In the H3_nuc_, a hexasome with a single dimeric H2A/H2B can maintain the integrity of a single DNA wrap around the histones.

In Figure 8A, it can be seen that there is an H3_nuc_(green arrow) that is wrapped at the 601 location (frame A1). In the following frames (A2 and A20), the histone (blue arrow) has vacated the octameric core, but the partial core remains to wrap the DNA. In frame A36, the DNA unwraps the partial core, resulting in a wrapping of ∼100 bp, and the free histone remains near. In the last frame, A66, the only histone remaining appears to be the tetramer. These predictions on the dimer being the first to leave the octameric core are based on the size of the histone leaving, the location, and the assembly process of nucleosomes, showing that dimers are the last ones to leave; it would make sense that they would be the first histones to leave.

H3_nuc_(green arrow), in Figure 8B, starts fully wrapped at the 601 location (frame B1). In frames B4 and B7, it can be seen that a histone (blue arrow) left the core particle and now drifts nearby. During this process, unwrapping of the DNA took the ∼150 bp wrapping down to ∼100 bp in frame B4. This unwrapping continues, and by frame B11, the core particle is no longer wrapping any DNA. We assume that the octameric core splits into its three components: H2A/H2B dimers (blue arrows) and H3/H4 tetramers (yellow arrow).

The CENP-A_nuc_, conversely to the H3nuc, had a much lower occurrence of the histones vacating the octameric core. In Figure 9A, the CENP-A_nuc_ (green arrow) can be seen sitting at the 601 location in frames A1, A4 and A18. In frame B20, the DNA on the short arm can be seen to have lengthened, indicating an unwrapping of the nucleosome (yellow arm now). Of note, no histones left the nucleosome during this unwrapping transition. The under-wrapped nucleosome can be seen stably bound to the DNA in frame A37, still at the 601 location.

Another example of the CENP-A_nuc_, unwrapping without the loss of a histone, is seen in Figure 9B. In this example, the CENP-A_nuc_ is not bound to the 601 site but near the 3WJ, indicating that this process is not DNA sequence-dependent. In frames B3, B8, and B11, the CENP-A_nuc_ is fully wrapped and stably wrapping the DNA. In frame 13, the DNA is unwrapped from the short arm (3WJ arm), but once again, the nucleosome (yellow arrow) does not lose any histones in the process. The nucleosome is eventually evacuated from the DNA in frame 18 (yellow arrow).

These differences between the H3 and CENP-A nucleosomes indicate an intrinsic difference between the interactions of the DNA and the nucleosomes. Despite the lower wrapping efficiency of CENP-A nucleosomes compared with H3_nuc_, we have shown here that the nucleosomes not only have longer dwell times on the DNA, but their octameric structural integrity is greater than that of H3_nuc_.

We also found that a step-wise disassembly process occurs in both H3_nuc_ and CENP-A_nuc_, resulting in a stable under-wrapped state of ∼110 bp. This step-wise disassembly is only unwrapped from a single side, as seen in Figures 10 and 11. Therefore, these nucleosomes had an entry/exit DNA that remains completely intact, while the other unraveled between 20 and 40 bp. These results give insight into how the nucleosomes may be translocated in a rolling fashion, breaking only the contacts at a single entry/exit site. The exciting thing about this under-wrapped state is that it appears more stable than the fully wrapped state, with more time for both nucleosomes to be under-wrapped instead of the fully wrapped state.

The asymmetric unwrapping of nucleosomes is another property observed in both nucleosome types. The asymmetry is the preferential pathway for the nucleosome unwrapping observed in 87% of cases for H3_nuc_ and 88% of the CENP-A_nuc_. Previously, asymmetry was observed for the initial stage of unwrapping for the breathing of DNA.^44^ Other published work discovered that the histone dimers (H2A/H2B) are the first to leave the octameric core, guided by the asymmetrical unwrapping of the DNA.^45,46^ Here, we observed the asymmetry in the nucleosome unwrapping over the entire unraveling process. Also, we observed the asymmetry in unwrapping for CENP-A nucleosomes, where the core remains intact, suggesting that the core dissociation is not a factor contributing to the asymmetry of the nucleosome unwrapping.

Overall, a variety of unique structural characteristics of canonical and centromere nucleosomes at the nanoscale have been found. We found that CENP-A nucleosomes are more stable than canonical nucleosomes, regardless of their lower wrapping efficiency. Moreover, time-lapse experiments demonstrate that nucleosomes with ∼100 bp DNA wrapped are in a transient state with elevated stability. These findings suggest that the amount of DNA wrapped around the histone core is not the only factor defining the nucleosome stability, instead, other interactions between the histone cores and DNA contribute to the stability. The unwrapping process is highly asymmetric, and it was observed with both types of nucleosomes, revealing a novel property of the nucleosome dynamics. Additionally, HS-AFM revealed higher stability of CENP-A nucleosomes compared with H3 nucleosomes, in which dissociation of the histone core occurs prior to the H3 nucleosome dissociation. The histone core of CENP-A nucleosomes remains intact even after the dissociation of DNA, so the re-assembly of the CENP-A nucleosomes is facilitated. This feature of CENP-A nucleosomes can be important for the centromere dynamics during mitosis and chromatin replication.

## ASSOCIATED CONTENT

Additional analysis of nucleosome location and wrapping for high-speed AFM movies (DOC) and raw high-speed AFM movies (Movie S1-S9) are provided. (AVI)

## AUTHOR INFORMATION

### Author Contributions

Conceptualization, Y.L.L.; methodology, S.F. and Z.S.; validation, S.F. and Z.S.; formal analysis, S.F.; investigation, S.F.; resources, Y.L.L.; writing—original draft preparation, S.F.; writing—review and editing, S.F.; visualization, S.F.; supervision, Y.L.L.; project administration, Y.L.L.; funding acquisition, Y.L.L. All authors have read and agreed to the published version of the manuscript.

## ACKNOWLEDGMENT

We thank Dr. Lyubchenko lab members for useful insight.

## ABBREVIATIONS

3WJ: three-way junction
AFM: atomic force microscopy
H3nuc: CENP-Anuc
CENP-A: containing nucleosomes
H3: containing nucleosomes

## REFERENCES

(1) McGinty, R. K.; Tan, S. Nucleosome Structure and Function. Chem. Rev. 2015, 115 (6), 2255–2273. 10.1021/cr500373h.

(2) Onufriev, A. V.; Schiessel, H. The Nucleosome: From Structure to Function through Physics. Curr. Opin. Struct. Biol. 2019, 56, 119–130. 10.1016/j.sbi.2018.11.003.

(3) Zhou, K.; Gaullier, G.; Luger, K. Nucleosome Structure and Dynamics Are Coming of Age. Nat. Struct. Mol. Biol. 2019, 26 (1), 3–13. 10.1038/s41594-018-0166-x.

(4) Pan, Y. G.; Banerjee, S.; Zagorski, K.; Shlyakhtenko, L. S.; Kolomeisky, A. B.; Lyubchenko, Y. L. A Molecular Model of the Surface-Assisted Protein Aggregation Process; preprint; Biophysics, 2018. 10.1101/415703.

(5) Lyubchenko, Y. L. Atomic Force Microscopy Methods for DNA Analysis. In Encyclopedia of Analytical Chemistry; Meyers, R. A., Ed.; Wiley, 2019; pp 1–31. 10.1002/9780470027318.a9258.pub2.

(6) Clapier, C. R.; Cairns, B. R. The Biology of Chromatin Remodeling Complexes. Annu. Rev. Biochem. 2009, 78 (1), 273–304. 10.1146/annurev.biochem.77.062706.153223.

(7) Becker, P. B.; Workman, J. L. Nucleosome Remodeling and Epigenetics. Cold Spring Harb. Perspect. Biol. 2013, 5 (9), a017905–a017905. 10.1101/cshperspect.a017905.

(8) Forties, R. A.; North, J. A.; Javaid, S.; Tabbaa, O. P.; Fishel, R.; Poirier, M. G.; Bundschuh, R. A Quantitative Model of Nucleosome Dynamics. Nucleic Acids Res. 2011, 39 (19), 8306– 8313. 10.1093/nar/gkr422.

(9) Stormberg, T.; Stumme-Diers, M.; Lyubchenko, Y. L. Sequence-Dependent Nucleosome Nanoscale Structure Characterized by Atomic Force Microscopy. FASEB J. 2019, 33 (10), 10916–10923. 10.1096/fj.201901094R.

(10) Stormberg, T.; Vemulapalli, S.; Filliaux, S.; Lyubchenko, Y. L. Effect of Histone H4 Tail on Nucleosome Stability and Internucleosomal Interactions. Sci. Rep. 2021, 11 (1). 10.1038/s41598-021-03561-9.

(11) Stormberg, T.; Lyubchenko, Y. L. The Sequence Dependent Nanoscale Structure of CENP-A Nucleosomes. Int. J. Mol. Sci. 2022, 23 (19), 11385. 10.3390/ijms231911385.

(12) Sun, Z.; Stormberg, T.; Filliaux, S.; Lyubchenko, Y. L. Three-Way DNA Junction as an End Label for DNA in Atomic Force Microscopy Studies. Int. J. Mol. Sci. 2022, 23 (19), 11404. 10.3390/ijms231911404.

(13) Wang, Y.; Stormberg, T.; Hashemi, M.; Kolomeisky, A. B.; Lyubchenko, Y. L. Beyond Sequence: Internucleosomal Interactions Dominate Array Assembly. J. Phys. Chem. B 2022, 126 (51), 10813–10821. 10.1021/acs.jpcb.2c05321.

(14) Lancrey, A.; Joubert, A.; Duvernois-Berthet, E.; Routhier, E.; Raj, S.; Thierry, A.; Sigarteu, M.; Ponger, L.; Croquette, V.; Mozziconacci, J.; Boulé, J.-B. Nucleosome Positioning on Large Tandem DNA Repeats of the ‘601’ Sequence Engineered in Saccharomyces Cerevisiae. J. Mol. Biol. 2022, 434 (7), 167497. 10.1016/j.jmb.2022.167497.

(15) Cutter, A. R.; Hayes, J. J. A Brief Review of Nucleosome Structure. FEBS Lett. 2015, 589 (20PartA), 2914–2922. 10.1016/j.febslet.2015.05.016.

(16) McGhee, J. D.; Felsenfeld, G. Nucleosome Structure.

(17) Zhou, B.-R.; Yadav, K. N. S.; Borgnia, M.; Hong, J.; Cao, B.; Olins, A. L.; Olins, D. E.; Bai, Y.; Zhang, P. Atomic Resolution Cryo-EM Structure of a Native-like CENP-A Nucleosome Aided by an Antibody Fragment. Nat. Commun. 2019, 10 (1), 2301. 10.1038/s41467-019-10247-4.

(18) Ohtomo, H.; Kurita, J.; Sakuraba, S.; Li, Z.; Arimura, Y.; Wakamori, M.; Tsunaka, Y.; Umehara, T.; Kurumizaka, H.; Kono, H.; Nishimura, Y. The N-Terminal Tails of Histones H2A and H2B Adopt Two Distinct Conformations in the Nucleosome with Contact and Reduced Contact to DNA. J. Mol. Biol. 2021, 433 (15), 167110. 10.1016/j.jmb.2021.167110.

(19) Luger, K. Crystal Structure of the Nucleosome Core Particle at 2.8 Å Resolution. 1997, 389.

(20) Park, P. J. ChIP-Seq: Advantages and Challenges of a Maturing Technology. Nat. Rev. Genet. 2009, 10 (10), 669–680. 10.1038/nrg2641.

(21) Poirier, M. G.; Oh, E.; Tims, H. S.; Widom, J. Dynamics and Function of Compact Nucleosome Arrays. Nat. Struct. Mol. Biol. 2009, 16 (9), 938–944. 10.1038/nsmb.1650.

(22) Szerlong, H. J.; Hansen, J. C. Nucleosome Distribution and Linker DNA: Connecting Nuclear Function to Dynamic Chromatin structureThis Paper Is One of a Selection of Papers Published in a Special Issue Entitled 31st Annual International Asilomar Chromatin and Chromosomes Conference, and Has Undergone the Journal’s Usual Peer Review Process. Biochem. Cell Biol. 2011, 89 (1), 24–34. 10.1139/O10-139.

(23) McKinley, K. L.; Cheeseman, I. M. The Molecular Basis for Centromere Identity and Function. Nat. Rev. Mol. Cell Biol. 2016, 17 (1), 16–29. 10.1038/nrm.2015.5.

(24) Pidoux, A. L.; Allshire, R. C. The Role of Heterochromatin in Centromere Function. In Philosophical Transactions of the Royal Society B: Biological Sciences; Royal Society, 2005; Vol. 360, pp 569–579. 10.1098/rstb.2004.1611.

(25) Stellfox, M. E.; Bailey, A. O.; Foltz, D. R. Putting CENP-A in Its Place. Cell. Mol. Life Sci. 2013, 70 (3), 387–406. 10.1007/s00018-012-1048-8.

(26) Boopathi, R.; Danev, R.; Khoshouei, M.; Kale, S.; Nahata, S.; Ramos, L.; Angelov, D.; Dimitrov, S.; Hamiche, A.; Petosa, C.; Bednar, J. Phase-Plate Cryo-EM Structure of the Widom 601 CENP-A Nucleosome Core Particle Reveals Differential Flexibility of the DNA Ends. Nucleic Acids Res. 2020, 48 (10), 5735–5748. 10.1093/nar/gkaa246.

(27) Sekulic, N.; Bassett, E. A.; Rogers, D. J.; Black, B. E. The Structure of (CENP-A-H4) 2 Reveals Physical Features That Mark Centromeres. Nature 2010, 467 (7313), 347–351. 10.1038/nature09323.

(28) Stumme-Diers, M. P.; Banerjee, S.; Hashemi, M.; Sun, Z.; Lyubchenko, Y. L. Nanoscale Dynamics of Centromere Nucleosomes and the Critical Roles of CENP-A. Nucleic Acids Res. 2018, 46 (1), 94–103. 10.1093/nar/gkx933.

(29) Stumme-Diers, M. P.; Banerjee, S.; Sun, Z.; Lyubchenko, Y. L. Assembly of Centromere Chromatin for Characterization by High-Speed Time-Lapse Atomic Force Microscopy. In Nanoscale Imaging; Lyubchenko, Y. L., Ed.; Methods in Molecular Biology; Springer New York: New York, NY, 2018; Vol. 1814, pp 225–242. 10.1007/978-1-4939-8591-3_14.

(30) Stumme-Diers, M. P.; Sun, Z.; Lyubchenko, Y. L. Probing the Structure and Dynamics of Nucleosomes Using AFM Imaging. 2020.

(31) Stormberg, T.; Filliaux, S.; Baughman, H. E. R.; Komives, E. A.; Lyubchenko, Y. L. Transcription Factor NF-κB Unravels Nucleosomes. Biochim. Biophys. Acta - Gen. Subj. 2021, 1865 (9), 129934. 10.1016/j.bbagen.2021.129934.

(32) Vemulapalli, S.; Hashemi, M.; Lyubchenko, Y. L. Site-Search Process for Synaptic Protein-Dna Complexes. Int. J. Mol. Sci. 2022, 23 (1). 10.3390/ijms23010212.

(33) Vemulapalli, S.; Hashemi, M.; Kolomeisky, A. B.; Lyubchenko, Y. L. DNA Looping Mediated by Site-Specific SfiI–DNA Interactions. J. Phys. Chem. B 2021, 125 (18), 4645–4653. 10.1021/acs.jpcb.1c00763.

(34) Miyagi, A.; Ando, T.; Lyubchenko, Y. L. Dynamics of Nucleosomes Assessed with Time-Lapse High-Speed Atomic Force Microscopy. Biochemistry 2011, 50 (37), 7901–7908. 10.1021/bi200946z.

(35) Lowary, P. T.; Widom, J. Nucleosome Packaging and Nucleosome Positioning of Genomic DNA (Chromatingene Regulationtranscriptional Activation); 1997; Vol. 94, pp 1183–1188. www.pnas.org.

(36) Widom, J. Role of DNA Sequence in Nucleosome Stability and Dynamics. Q. Rev. Biophys. 2001, 34 (3), 269–324. 10.1017/S0033583501003699.

(37) Menshikova, I.; Menshikov, E.; Filenko, N.; Lyubchenko, and Y. L. Nucleosomes Structure and Dynamics: Effect of CHAPS. www.ijbmb.org.

(38) Lyubchenko, Y. L. Direct AFM Visualization of the Nanoscale Dynamics of Biomolecular Complexes. J. Phys. Appl. Phys. 2018, 51 (40), 403001. 10.1088/1361-6463/aad898.

(39) Lyubchenko, Y. L.; Shlyakhtenko, L. S. Chromatin Imaging with Time-Lapse Atomic Force Microscopy. In Chromatin Protocols; Chellappan, S. P., Ed.; Methods in Molecular Biology; Springer New York: New York, NY, 2015; Vol. 1288, pp 27–42. 10.1007/978-1-4939-2474-5_3.

(40) Lyubchenko, Y. L. Nanoscale Nucleosome Dynamics Assessed with Time-Lapse AFM. Biophys. Rev. 2014, 6 (2), 181–190. 10.1007/s12551-013-0121-3.

(41) Lyubchenko, Y. L.; Shlyakhtenko, L. S.; Ando, T. Imaging of Nucleic Acids with Atomic Force Microscopy. Methods 2011, 54 (2), 274–283. 10.1016/j.ymeth.2011.02.001.

(42) Chua, E. Y. D.; Vasudevan, D.; Davey, G. E.; Wu, B.; Davey, C. A. The Mechanics behind DNA Sequence-Dependent Properties of the Nucleosome. Nucleic Acids Res. 2012, 40 (13), 6338–6352. 10.1093/nar/gks261.

(43) Kixmoeller, K.; Allu, P. K.; Black, B. E. The Centromere Comes into Focus: From CENP-A Nucleosomes to Kinetochore Connections with the Spindle. Open Biol. 2020, 10 (6), 200051. 10.1098/rsob.200051.

(44) Ngo, T. T. M.; Ha, T. Nucleosomes Undergo Slow Spontaneous Gaping. Nucleic Acids Res. 2015, 43 (8), 3964–3971. 10.1093/nar/gkv276.

(45) Chen, Y.; Tokuda, J. M.; Topping, T.; Meisburger, S. P.; Pabit, S. A.; Gloss, L. M.; Pollack, L. Asymmetric Unwrapping of Nucleosomal DNA Propagates Asymmetric Opening and Dissociation of the Histone Core. Proc. Natl. Acad. Sci. 2017, 114 (2), 334–339. 10.1073/pnas.1611118114.

(46) Zhang, B.; Zheng, W.; Papoian, G. A.; Wolynes, P. G. Exploring the Free Energy Landscape of Nucleosomes. J. Am. Chem. Soc. 2016, 138 (26), 8126–8133. 10.1021/jacs.6b02893.

