## Supplemental data for "Nanoscale Structure, Interactions, and Dynamics of Centromere Nucleosomes"

### Supplementary Figures

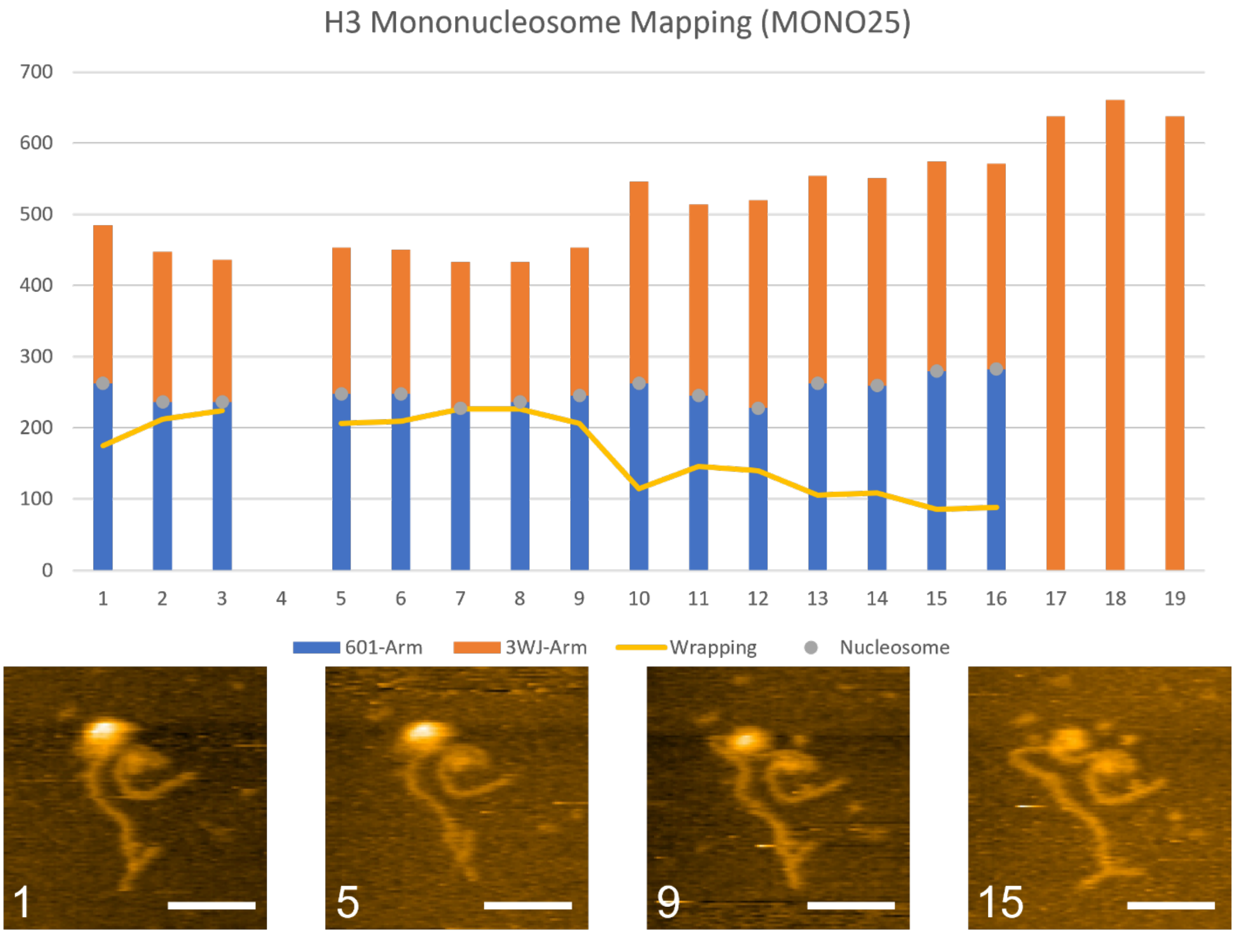


**Supplementary Fig. S1. HS-AFM analysis of $H3_{nuc}$.** The Movie S5 video analyzed, showing the mapping (bar graph) and snap shots of particular frames. The grey dots in the bar graph indicate the nucleosome bound to the DNA. The yellow line shows the wrapping of the $H3_{nuc}$. The number in white in the bottom left of the snap shots indicates the frame the image is from.

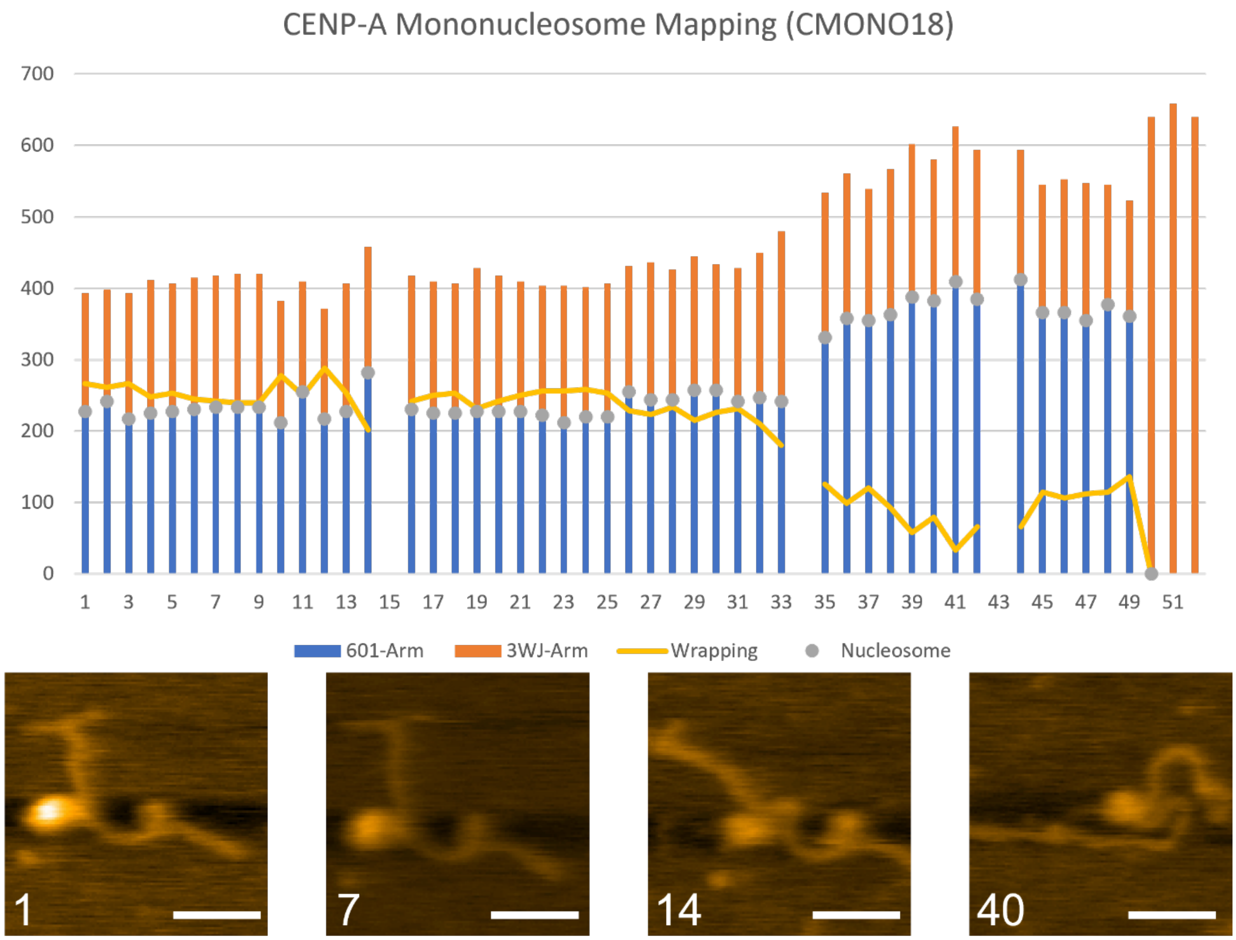


**Supplementary Fig. S2. HS-AFM analysis of CENP-A$_{nuc}$.** The Movie S9 video analyzed, showing the mapping (bar graph) and snap shots of particular frames. The grey dots in the bar graph indicate the nucleosome bound to the DNA. The yellow line shows the wrapping of the CENP-A$_{nuc}$. The number in white in the bottom left of the snap shots indicates the frame the image is from.
